# Hypothesis Tests for Neyman’s Bias in Case-Control Studies

**DOI:** 10.1101/066902

**Authors:** D.M. Swanson, C.D. Anderson, R.A. Betensky

## Abstract

Survival bias is a long-recognized problem in case-control studies, and many varieties of bias can come under this umbrella term. We focus on one of them, termed Neyman’s bias or “prevalence-incidence bias.” It occurs in case-control studies when exposure affects both disease and disease-induced mortality, and we give a formula for the observed, biased odds ratio under such conditions. We compare our result with previous investigations into this phenomenon and consider models under which this bias may or may not be important. Finally, we propose three hypothesis tests to identify when Neyman’s bias may be present in case-control studies. We apply these tests to three data sets, one of stroke mortality, another of brain tumors, and the last of atrial fibrillation, and find some evidence of Neyman’s bias in the former two cases, but not the last case.

## Introduction

Survival bias is a frequent source of concern in case-control studies (Sackett, 1979; Rothman et al., 2008). Sackett describes nine types of bias common in case-control studies, and we focus our investigation on one of them, first identified by Jerzy Neyman and now known as Neyman’s bias or “prevalence-incidence bias” (Neyman, 1955). It is a bias that occurs when prevalent cases are sampled and exposure affects disease and disease-associated mortality. Since Neyman’s article was written in the 1950’s when the relationship between smoking and lung cancer was under debate, he uses an example that focuses on that subject. He disregards competing risks and supposes that if, in fact, smoking is protective against lung cancer, but lung cancer mortality is far higher among non-smokers than smokers, then the odds ratio would suggest that smoking is a risk factor for disease as was being observed at the time. In our study of the subject, we focus on three other examples, one coming from a study of brain tumors and chemotherapy, another coming from a GWAS of ischemic stroke, and the last coming from a study of atrial fibrillation in the Framingham Heart Study. Prevalence-incidence bias could arise in the study of brain tumors if certain patients are assessed to have disease too progressed to benefit from chemotherapy and therefore do not undergo treatment. The GWAS could suffer from prevalence-incidence bias if a certain subset of patients die before admission to a hospital and study entry. We use our study of atrial fibrillation in the Framingham Heart study as an example of a prospective design that should therefore not suffer from prevalence-incidence bias. We consider these data as being generated under the null hypothesis of no prevalence-incidence bias in order to substantiate the validity of the test.

Despite Neyman’s early identification of this bias, methodological investigation into it has been limited. Hill (2003) uses a compartment model to show how bias arises when performing case-control studies on prevalent cases if the risk factor impacts both disease and mortality from disease. He also shows that any impact of the risk factor on mortality from other causes does not impact the observed odds ratio, which demonstrates that Neyman was justified in ignoring competing risks. While trying to draw inference on incidence instead of the odds ratio, Fluss et al. (2012), Keiding (1991), and Keiding (2006) all consider the problem of using cross-sectional designs and their resultant sampling biases.

Anderson et al. (2011) performs a computational investigation into Neyman’s bias, recognizing that genome-wide association studies (GWAS) and their use of prevalent cases in case-control study designs were susceptible to it. If an allele is a risk factor for both disease and mortality from disease, then the common practice of calculating an odds ratio from prevalent cases and controls could lead to biased inference. Since the odds ratios in such studies are usually small, differences in disease-associated mortality between the exposed and unexposed would not be required for a risk allele to be observed as protective, or vice versa. Their own investigation is motivated by a locus found to be significantly associated with ischemic stroke in longitudinal studies that did not replicate using a case-control design. As a solution, they simulate data under different disease and mortality risk models and then fit regression models for percent bias of the odds ratio to the disease and mortality risk model parameters. These fitted models give researchers a means to investigate the potential biases of estimated odds ratios in their own studies.

In this paper, we propose a framework for consideration of Neyman’s bias and examine it from a modeling perspective. We suggest three hypothesis tests to assess whether Neyman’s bias is present in a study and then apply these tests to three data sets mentioned: one of brain tumors and chemotherapy, another a GWAS of ischemic stroke, and the last focused on atrial fibrillation in the Framingham Heart Study. We propose hypothesis tests rather than methods to recover the true, unbiased odds ratio, since that quantity is unrecoverable under the framework we consider.

## Methods

### 2.1 Notation and background

Suppose that we have a setting similar to that described in Anderson et al. (2011), where we have some binary risk SNP or gene, G, that takes on the value 1 with probability *p* (“exposed”) and 0 with probability 1 – *p* (“unexposed”). Let *D* denote age of disease onset, and suppose that *G* may be associated with *D*. Let {*M_a,j_*}, *j* = 1,…, *n*, denote age at mortality from all other causes not associated with disease. Let {*X_i_*}, *i* = 1,…, *m*, denote latent time from disease onset to the *i^th^* mortality cause related to disease and let *X* = min{*X_i_*}. Thus, *M_a,j_* ╨ (*X D*)^*T*^ | *G* for all *j*, where *W* ╨ *Y* | *Z* denotes statistical independence of *W* and *Y* conditional on *Z* (Dawid, 1979). We define *M_a_* ≡ min{*M_a,j_*}, and thus *M_a_* ╨ (*X D*)^*T*^ | *G*. Let *M_d,i_* ≡ *D* + *X_i_* and *M_d_* = min(*M_d,i_*) = *D* + *X*. If *X_i_* ╨ *D*, then *M_d,i_* is necessarily associated with *D* because *M_d,i_* ≡ *D* + *X_i_* (i.e., *M_d,i_* denotes the age at disease-associated mortality cause *i*). In fact, *X_i_* would have to be associated with *D* in a specific way to have *M_d,i_* ╨ *D*. We do not assume *X_i_* is a positive random variable so that we can have *P*(*M_d,i_* < *D*) ≥ 0. While it may seem counterintuitive to allow for disease-associated mortality prior to disease, this flexibility fits into a realistic framework. For example, if the disease of interest is stroke, and there exists an association between death from myocardial infarction and stroke, then indeed mortality associated with disease, though not directly caused by it, can occur before disease and can bias the odds ratio, as we show later.

It is not a limitation of this conceptual framework to assume the existences of the *M_a,j_*’s, ages at causes of mortality unrelated to our disease of interest. They are present to show their lack of effect on the observed odds ratio in the work to follow. Since there is no cap on the possible number of *M_d,i_*’s, all causes of mortality can be considered as disease-associated if desired by the analyst.

### 2.2 Formulae

Suppose we perform a case-control study of prevalent cases at age *t*\*, and define *C_a_* ≡ *I*(*t*\* ≤ *M_a_*), *C_d_* ≡ *I*(*t*\* ≤ *M_d_*), where *I*(·) is the indicator function, and *C* ≡ *C_d_* × *C_a_*. While *C_d_* and *C_a_* are functions of *M_d_* and *M_a_*, mortality causes, we can consider *M_d_* and *M_a_* more generally as anything that would render a subject unable to enter the study that is associated and unassociated with disease, respectively. A subject is available to enter the study at age *t*\* if *C* = 1; i.e., if the subject has not died from any cause by age *t*\*. Denote the cumulative distribution function associated with random variable *Y* as *F_Y_* (*t*). Then the target odds ratio among the population at age *t*\* is

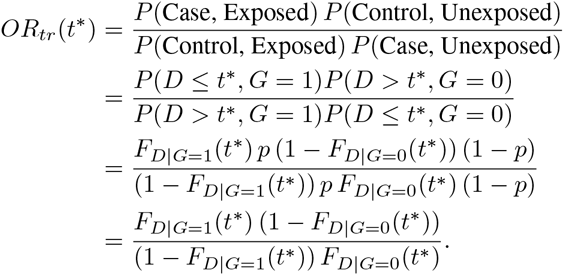

In contrast, the observed odds ratio among prevalent cases at age *t*\* iss

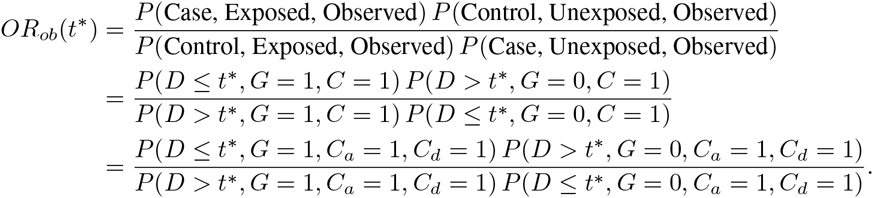

Now consider the term *P* (*D* ≤ *t*\*, *G* = 1, *C_a_* = 1, *C_d_* = 1). We can factor the probability as

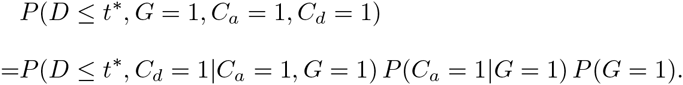

Since *M_a_* ╨ (*X D*)^*T*^ | *G* and *M_d_* ≡ *X* + *D, M_a_* ╨ *M_d_* | *G*, and since *C_a_* and *C_d_* are functions of only *M_a_* and *M_d_* (with fixed and known *t*\*), respectively, (*D C_d_*)^*T*^ ╨ *C_a_* | *G*. Using this conditional independence, the probability further simplifies to

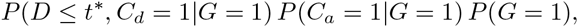

which is equal to

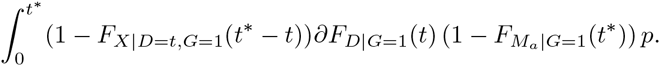

Analogous simplifications of the other terms of *OR_ob_*(*t*\*) yield

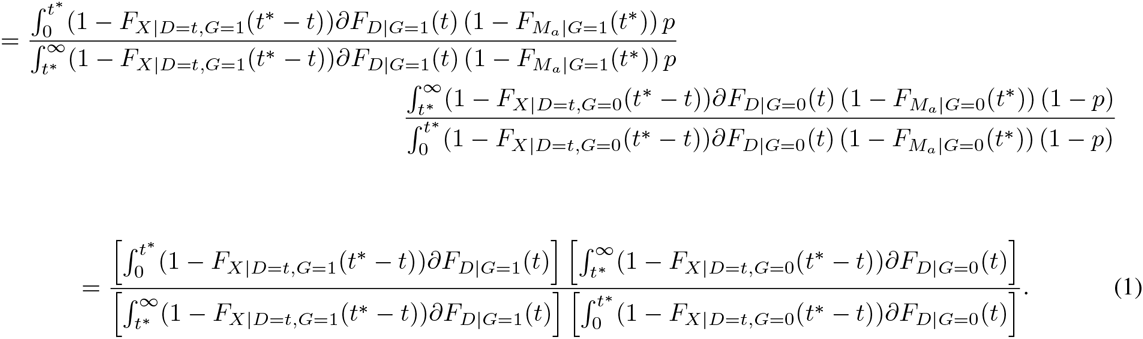

Note that when *X* ╨ *D|G*, we observe

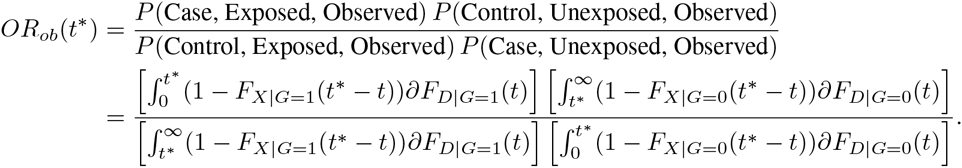

This assumption may be reasonable for some exposures that are risk factors for diseases whose course is independent of the age of onset given *G*.

Returning to the general case (1), we consider ways in which *OR_ob_*(*t*\*) = *OR_tr_*(*t*\*) holds. Recall that *M_d_* ≡ *D* + *X*, and that *X* need not be a positive random variable. Suppose that *X* ≡ *A − D*, for some positive random variable *A* independent of *D*, conditional on *G*. Then *M_d_* ≡ *D* + *X* = *D* + (*A − D*) = *A*. So *M_d_* = *A* and is independent of *D* given *G*, or in notation, *M_d_* ╨ *D* | *G* (Dawid, 1979). Notice that when *M_d_* is defined in this way, an association necessarily exists between *X* and *D*, conditional on *G*, since *X* is itself a function of *D*. If *M_d_* ╨ *D* | *G* holds, then (1) reduces to

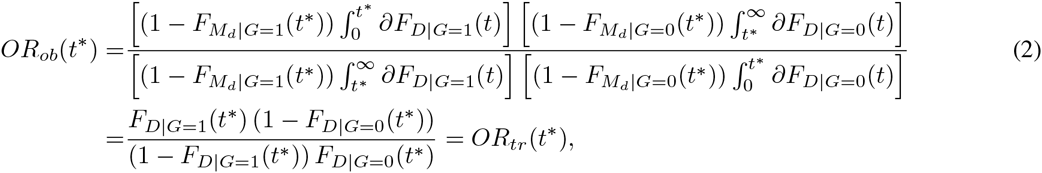

where (2) follows from

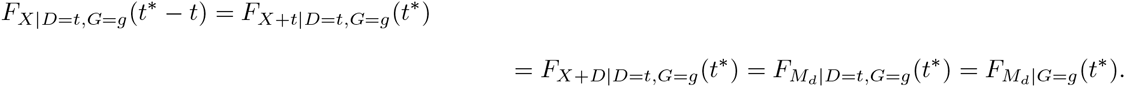

So when *M_d_* ╨ *D | G, M_d_* behaves like *M_a_* in the sense that *OR_ob_*(*t*\*) is no longer a function of the distribution of *M_d_* and *OR_tr_*(*t*\*) = *OR_ob_*(*t*\*). While *M_d_* ╨ *D* | *G* is a sufficient condition for *OR_tr_*(*t*\*) = *OR_ob_*(*t*\*), it is not necessary; there exist multivariate distributions (*X D G*)^*T*^ such that *OR_tr_*(*t*\*) = *OR_ob_*(*t*\*), but 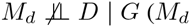 is not independent of *D* conditional on *G*). For example, consider the case in which *F_M_d_|D=t,G=g_*(*x*) =0 if *x* ≤ *t*\* or *x* ≤ *t* and *F_M_d_|D=t,G=g_*(*x*) = 1, otherwise for *g* ∈ {0,1}; i.e., no disease-related death occurs prior to *t*\* and in this way cannot bias *OR_ob_*(*t*\*), but in the region *D* > *t*\*, *M_d_* is perfectly correlated with *D* so that 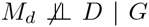. Nonetheless, in § 5 we propose tests of deviations from *M_d_* ╨ *D* | *G* since the cases in which 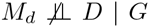, but *OR_tr_*(*t*\*) = *OR_ob_*(*t*\*) holds are unlikely to occur as is the case in this example.

## Scientific hypotheses versus sampling bias hypotheses

We distinguish between *H*_0*S*_: *OR_tr_*(*t*\*) = 1 (at some time *t*\*, the true odds ratio is one), which we term the “scientific null hypothesis” and *H*_0*B*_: *OR_tr_*(*t*\*) = *OR_ob_*(*t*\*) (there is no bias in the odds ratios at time *t*\*), which we term the “sampling bias null hypothesis.” The alternative hypothesis in each case is the complement of the null hypothesis. We describe characteristics of these hypotheses.

*Under H*_0*S*_: *OR_tr_*(*t*\*) = 1 *and* 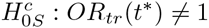:

Even if mortality from other causes, *M_a_*, depends on *G*, it does not affect the bias of the observed odds ratio; in other words, *OR_ob_*(*t*\*) and *OR_tr_*(*t*\*) are not a function of the distribution of *M_a_*. Thus, we may assume, as Neyman (1955) does in his original example and Hill (2003) confirms, that mortality from other causes is not present and death can only occur from disease. Similarly, the probability of exposure, *p*, does not affect *OR_ob_*(*t*\*). Also, if *F*_*M*_*d*_|*G*=*g*_ (*t*\*) = 0 for *g* ∈ {0,1} (which is the case when no disease-associated mortality occurs prior to *t*\*), then *OR_ob_*(*t*\*) is unbiased: *OR_ob_*(*t*\*) = *OR_tr_*(*t*\*). This result is expected since it is disease-related mortality that results in the bias-inducing differential selection between the exposed and unexposed.

*Under* 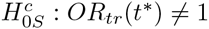:

Under the following four conditions, bias exists (i.e., *OR_ob_*(*t*\*) ≠ *OR_tr_*(*t*\*)):

1. 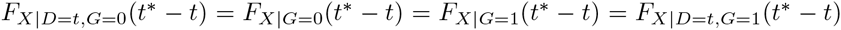 for all *t* (i.e., the mortality distribution from disease-onset is identical between the exposed and unexposed and not dependent on age at disease-onset).
2. *F_X_*_|*G=g*_(*t*^**^) > 0 for some *g* ∈ {0,1}. In other words, either the exposed or unexposed have positive probability of dying from disease by *t*^**^, where *t*^**^ is defined as the time between *t*\* and the first possible presence of disease among the exposed or unexposed (i.e., inf{*F*_*D*|*G*=*g*_(*t*) > 0: *t* ∈ [0, ∞), *g* ∈ {0,1}}) so that the bias-inducing event will have some chance of occurring prior to study entry at age *t*\*).
3. *P*(*X* > 0) = 1, implying *P*(*D < M_d_*) = 1.
4. *F*_*D*|*G*=0_(*x*) = *F*_*D*|*G*=1_(*x−k*) forall *x* for some *k* ≠ 0, and *F*_*D*|*G*=0_(*t*\*) > 0 or *F*_*D*|*G*=1_(*t*\*) > 0 (i.e., the disease distributions for the exposed and unexposed are in the same location family, and *k* ≠ 0 implies *OR_tr_*(*t*\*) = 1).

These assumptions seem plausible if some exposure affects the mean age of disease, though the shape of the disease distribution is approximately the same between exposed and unexposed, and after disease occurrence, hazard of mortality is identical among those with and without the exposure and not a function of age at disease onset. The theorem and proof of this result is found in the Appendix (Theorem 1). Additionally, in that proof we examine the direction of bias; we find that when *OR_tr_*(*t*\*) < 1, then *OR_ob_*(*t*\*) > *OR_tr_* (*t*\*), and when *OR_tr_*(*t*\*) > 1, then *OR_ob_*(*t*\*) < *OR_tr_*(*t*\*). Thus, if the degree of bias is relatively small, then it can be viewed as a bias toward an observed odds ratio of 1. However, *OR_ob_*(*t*\*) is by no means bounded by 1 and so if the amount of bias is great, *OR_ob_*(*t*\*) and *OR_tr_*(*t*\*) can lie on opposite sides of 1, leading to wrongly inferring a truly protective exposure as a risk factor for the outcome or a true risk factor as protective against the outcome.

This result of *OR_ob_*(*t*\*) ≠ *OR_tr_*(*t*\*) will not necessarily hold if conditions 1 – 3 hold, but condition 4 is not satisfied (the distributions of disease of exposed and unexposed are not in the same location family). Under such a scenario, there may not be bias, as Example 1 in the Appendix illustrates. Additionally, if we only assume that conditions 2 – 3 are satisfied, then there may or may not be bias. See Examples 2 and 3 in the Appendix for instances of *OR_ob_*(*t*\*) = *OR_tr_*(*t*\*) and *OR_ob_*(*t*\*) ≠ *OR_tr_*(*t*\*), respectively, when *X* is associated with *G* (but is independent of *D* given *G: X* ╨ *D* | *G*). It follows that if there exist no conditional independences, one can make no conclusions regarding the relationship between *OR_tr_*(*t*\*) and *OR_ob_*(*t*\*) as there is even greater flexibility in the joint model. Lastly, if only *X* ╨ *G | D* is assumed so that *X* may depend on *D* (i.e., time to disease-induced mortality may depend on age at disease-onset), again *OR_tr_*(*t*\*) and *OR_ob_*(*t*\*) may or may not be equal. This result follows from the proof with location families and Example 1 because they are special cases of only assuming *X ╨ G | D*.

*Under H*_0*S*_: *OR_tr_*(*t*\*) = 1:

If we only assume that *OR_tr_*(*t*\*) = 1 with no conditions on *OR_tr_*(*t*) for *t* < *t*\*, and also that *X* ╨ *D* | *G* and *F_X|G=0_*(*t*) ≠ *F_X|G=1_*(*t*) for some *t* < *t*^**^, one cannot conclude anything regarding the relationship between *OR_tr_*(*t*\*) and *OR_ob_*(*t*\*). Consider Examples 4 and 5 in the Appendix for instances of *OR_ob_*(*t*\*) = *OR_tr_*(*t*\*) = 1 and *OR_ob_*(*t*\*) ≠ *OR_tr_*(*t*\*) = 1, respectively. We also observe that if *OR_tr_*(*t*) = 1 for all *t* ≤ *t*\* and *F_X|D,G=0_*(*t*) = *F_X|D,G=1_*(*t*) for all *t* ≤ *t*^**^, *OR_tr_*(*t*\*) = *OR_ob_*(*t*\*) = 1.

## The odds ratio when *T*\* is not fixed

If the case-control study consists of people of many ages, then *t*\*, previously considered fixed, can be considered random. Let us denote this random variable *T*\*. Under these conditions, the target odds ratio becomes

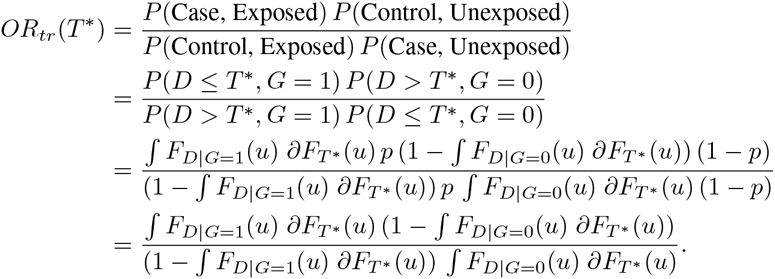

Making no assumptions about the joint model (*D X M_a_ G*)^*T*^, the observed odds ratio is

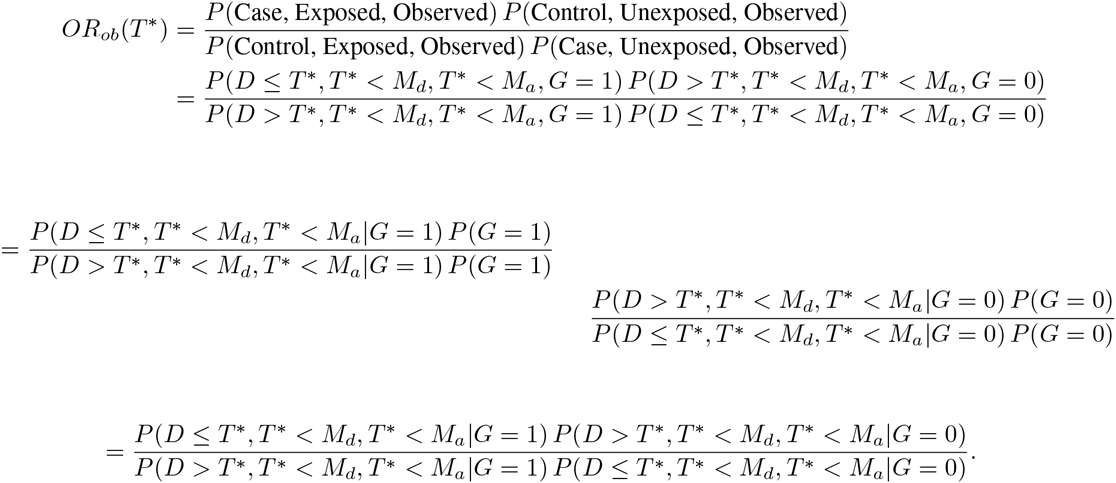

While *P*(*G* = 1) = *p* cancels from *OR_ob_*(*T*\*) as before with *OR_ob_*(*t*\*), we see that even if (*D X*)^*T*^ ╨ *M_a_|G*, we cannot factor *P*(*T*\* < *M_a_|G = g*) out of the expression. So *OR_ob_*(*T*\*) becomes a function of *M_a_*, causes of mortality unassociated with the disease under investigation. Additionally, regardless of whether *P*(*T*\* < *M_a_|G = g*) factors out of the expression, *D* ╨ *M_d_ | G*, which we have stated before as being sufficient for *OR_ob_*(*t*\*) = *OR_tr_*(*t*\*), is not sufficient for *OR_ob_*(*T*\*) = *OR_tr_*(*T*\*). This point is relevant as we propose hypothesis tests below. This is also important to remember since *OR_ob_*(*T*\*) is generally what would be measured in a real-world case-control study where ages of subjects vary. While outside the scope of this investigation, investigators might want to only pool those groups of subjects where *F*_*D*|*G*_(·) is relatively constant across their age ranges, in which case *D* ╨ *M_d | G_* would be sufficient for *OR_ob_*(*T*\*) = *OR_tr_*(*T*\*). Also, if the sample size of a case-control study is sufficient, stratifying subjects by age and calculating age-specific odds ratios would be another way to be assured that *D* ╨ *M_d_ | G* is sufficient for those stratum-specific odds ratios.

## Hypothesis testing

### 5.1 Description

We develop three methods for testing for the presence of Neyman’s bias in a study. Again, the “bias null hypothesis” of these tests is *OR_tr_*(*t*\*) = *OR_ob_*(*t*\*), and the alternative is *OR_tr_*(*t*\*) ≠ *OR_ob_*(*t*\*). While power may vary as a function of *OR_tr_*(*t*\*), the tests we propose are valid under all values of *OR_tr_*(*t*\*). Each of these three methods makes use of characteristics unique to the data when Neyman’s bias is absent, and each test may be more fitting to use than the other two under certain study designs. So, for example, Tests 1 and 2 require study observations to have some variation in age at study entry, a random variable we denote *T*\*, while Test 3 does not, though Test 3 requires external knowledge of population prevalence of disease and exposure, while neither Test 1 nor Test 2 does.

We have demonstrated above that *M_d_* ╨ *D* | *G* is a sufficient condition for *OR_tr_*(*t*\*) = *OR_ob_*(*t*\*). Ideally, we would have data on all of *D, M_d_*, and *G* and could test for conditional independences. However, in practice, it may be unlikely that one would have follow-up data on controls and perhaps even cases, in which case *M_d_* would be unknown for one or both groups. Thus, we propose these tests with real-world data limitations in mind.

The first two hypothesis tests we propose attempt to test whether this independence condition holds. Both of these hypothesis tests make use of previous work coming from the truncation methodology literature for tests of “quasi-independence,” which refers to independence of random variables in a certain “observable” region of their joint distribution, which we explain further below (Martin and Betensky, 2005; Tsai, 1990). Tests for quasi-independence are based on U-statistics, a class of statistics with broad application outside of these tests and first described in Hoeffding etal. (1948).

The last hypothesis test we propose assumes *P*(*D < M_d_*) = 1, which may be unreasonable in some settings, but reasonable in others, and depends on whether causes of mortality associated with disease can come before disease onset. The test uses the fact that with data collected under a case-control study design along with population disease prevalence, one can estimate the population exposure proportion. If one has knowledge of the true exposure proportion, any comparison between the true, known value and the calculated quantity can reveal bias in the odds ratio from which it was calculated. Thus, in contrast to the first two tests that detect a sufficient, though not necessary, condition for *OR_tr_*(*t*\*) = *OR_ob_*(*t*\*), rendering the test potentially slightly conservative (though likely not very conservative), this last test achieves its nominal type 1 error rate under the null of *OR_tr_*(*t*\*) = *OR_ob_*(*t*\*) and has power greater than it under the alternative of *OR_tr_*(*t*\*) ≠ *OR_ob_*(*t*\*).

### 5.2 Test 1: Testing for “quasi-independence” using *D* and *M_d_*

We are interested in testing independence of *D* and *M_d_* given *G*, and our observable region is *D* < *T*\* < *M_d_* given *G*; i. e., realizations of observed (because *T*\* < *M_d_*) cases (because *D* < *T*\*) of a given exposure status. To accomplish this, we modify a *U*-statistic test of association of Austin et al. (2013), whose null hypothesis assumes in our context mutual independence of *D, T*\*, *M_d_*; this is stronger than our null hypothesis. This is a valid approach to testing *D* ╨ *M_d_* | *G*, which is sufficient for no Neyman’s bias, because *D* ╨ *M_d_* | *G* necessarily implies independence in the region we are defining as observable, *D* < *T*\* < *M_d_* given *G*. Additionally, we focus on the cases in the study, under the assumption that follow-up data on Md is more likely to be available among them. While the power of this test may suffer in comparison to one that makes use of all observations, the approach makes fewer assumptions on data availability, and in settings in which *P*(*D* < *M_d_*) is close to 1, power will not suffer significantly.

To implement the hypothesis test, first we categorize all causes of mortality as *M_d_*, since if *D* and *M_d_* are associated given *G*, and *D* ╨ *M_a_* | *G*, then categorizing *M_a_* as *M_d_* will maintain that association and avoid the need to censor observations. Doing so is not an approximation nor does it invalidate the test; rather, the test could become invalid if mortality related to disease (*M_d_*) are incorrectly categorized as unrelated to disease (*M_a_*). Also, if *D* ╨ *M_d_* | G and *D* ╨ *M_a_* | *G*, categorizing *M_a_* as *M_d_* will maintain *D* ╨ *M_d_* | *G*. This approach is also legitimate from the perspective that *M_d_* was originally defined as causes of mortality potentially, though not necessarily, associated with disease. Now suppose that we have 1,…,*n* realizations of 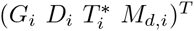, all cases so that one can assume *D* < *T*\* and on whom there is follow-up so *M_d_* is known, and that 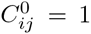 (alternatively, 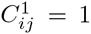) if *G* = 0 (alternatively, *G* =1) and max{*D_i_, D_j_*} ≤ min{*M_d,i_, M_d,j_*}, the *comparability* criterion, is satisfied, and 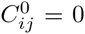 (altenatively, 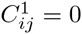) otherwise. Define 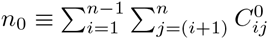 and 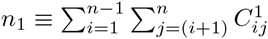.

The test statistic for the stratum *G* = *g, T_g_*, with *g* ∈ {0,1}, is

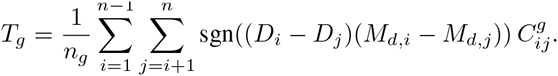

Then *T_g_* ~ *N*(0, *υ_g_*), where

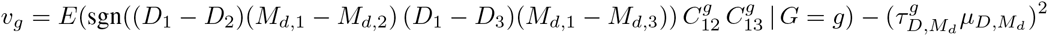

and where 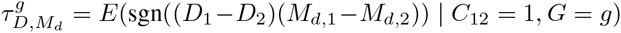 and *μ_D,M_d__* = *P*(*C*_12_ = 1), with sgn(*x*) = 1 for *x* > 0, − 1 for *x* < 0, and 0 for *x* = 0.

Since we would reject if either *T*_0_ or *T*_1_ falls in some predetermined critical region because dependence between *D* and *M_d_* given either *G* = 0 or *G* = 1 may mean *OR_tr_*(*t*\*) ≠ *OR_ob_*(*t*\*), in order to achieve a size *α* test, we can use a p-value threshold of *α*\* for *T*_0_ and *T*_1_, where *α*\* satisfies the equation *α* = 1 − (1 − *α*\*)^2^. So we propose a test that rejects for 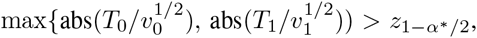 where abs(*x*) denotes the absolute value of *x* and *z*_1−*α*\*/2_ is the (*z*_1−*α*\*/2_)^*th*^ quantile of a standard normal random variable.

Also, though *D* ╨ *M_d_ | G* characterizes a subset of situations for which *OR_tr_*(*t*\*) = *OR_ob_*(*t*\*), our test is likely not overly conservative. The majority of situations in which *OR_tr_*(*t*\*) = *OR_ob_*(*t*\*) holds result from *D* ╨ *M_d_ | G* being satisfied.

Power curves for Test 1 as a function of the association between *D* and *M_d_* are shown in Figs. 1 and 2. These curves were generated at 11 different values of Kendall’s *τ*, used as a measure of the association between *D* and *M_d_*. In our case a Kendall’s *τ* value of 0 corresponds to independence between *D* and *M_d_*, and the power of the test at that value demonstrates the desired type 1 error rate of 0.05. The power curve in Fig. 1 was generated using 3000 iterations at each value, while that in Fig. 2 was generated using 1000 iterations at each value, and power was estimated by averaging over these iterations. In Fig. 1, at each iteration the test statistic was calculated using 1000 comparable pairs (i.e., those pairs that satisfy the comparability criterion mentioned in the description of Test 1). In Fig. 2, the test statistic was calculated using approximately 670 comparable pairs at each iteration–a subset of the 1000 comparable pairs used for Test 2, described below, that satisfied Test 1’s more stringent comparability criterion. Fig. 2 also demonstrates a type 1 error rate of 0.05. *D*, *T*, and *M_d_* were all distributed normal with means of 5, 9, and 9, respectively, and standard deviations of 0.7.

**Figure 1:**
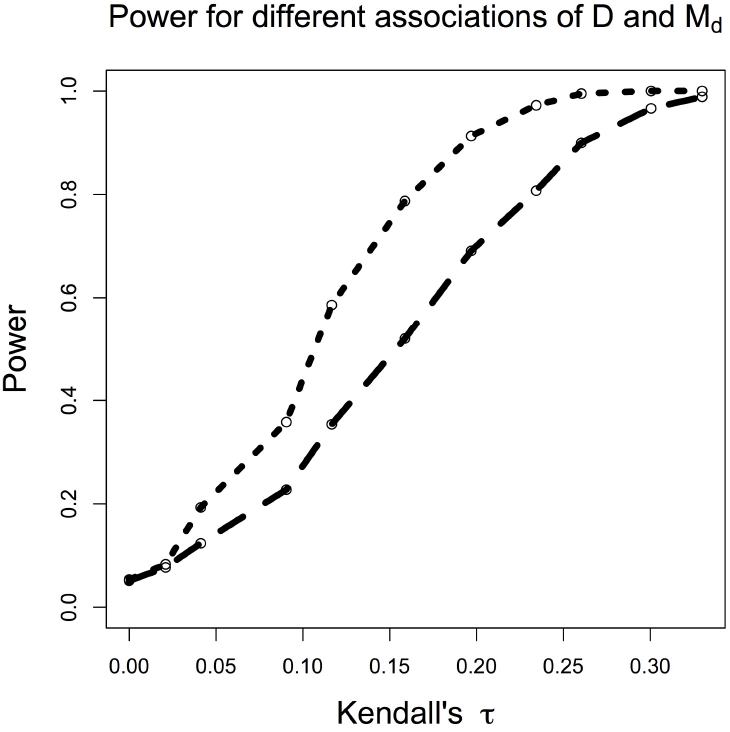
Comparison of power between tests 1 (short dashes) and 2 (long dashes) as a function of the association between *D* and *M_d_*, measured by Kendall’s *τ*, holding the sample size constant.

**Figure 2:**
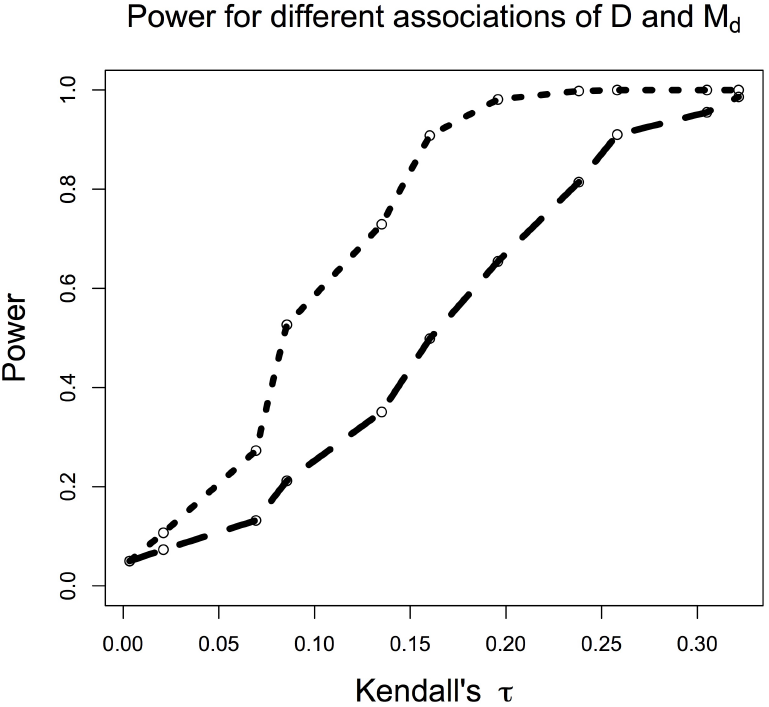
Comparison of power for tests 1 (short dashes) and 2 (long dashes) as a function of the *D* and *M_d_* as measured by Kendall’s *τ*, holding the number of comparable pairs constant.

We now consider realistic settings in which age of entry, *T*\*, is random. In § 4 above, we saw that the distribution for *M_a_* did not factor out of the odds ratio even when (*D X*) ╨ *M_a_ | G*, and that additionally even under the assumption of *D* ╨ *M_d_ | G*, whether or not the previous assumption held, *OR_ob_*(*t*\*) could be biased; we needed a fixed *t*\* for these conditional independencies to result in *OR_tr_*(*t*\*) = *OR_ob_*(*t*\*). Thus, it may seem illogical to propose a test that requires variation in *T*\*, which is precisely when the odds ratio will almost certainly be biased as shown in § 4. If we do find that *D* ╨ *M_d_ | G*, sufficient for no Neyman’s bias, we would need to then stratify our sample according to similar values of *T*\* such that, within each stratum, *T*\* can be effectively considered fixed, and then calculate the odds ratio for these different strata. We could then combine these strata into a average odds ratio if desirable or just consider each stratum-specific odds ratio separately.

### 5.3 Test 2: Testing “quasi-independence” with *D* and *t*\*

We now describe a test related to Test 1, which does not require knowledge of *M_d_* and again focuses on cases, those observations for whom *D < T*\* is true. Such a test is appropriate if a study did not obtain follow-up on subjects, but did record age at onset of disease for cases. The foundation for the test is based on causal directed acyclic graphs (DAGs), borrowed from the causal inference literature (Hernan and Robins, ress). We use DAGs not for the sake of justifying causal interpretations of *OR_ob_*, but rather as a convenient means of encoding conditional independencies. If DAGs are unfamiliar with the reader, Hernan and Robins (ress) describes them well.

By definition, the event *I*(*T*\* < *M_d_*) = 1 must be satisfied for any subject in the study and can therefore be treated as a conditioning event. Additionally, by definition of *I*(*T*\* < *M_d_*), there exists an association between it and both *T*\* and *M_d_*; to borrow language from the causal inference literature, *I*(*T*\* < *M_d_*) is called a “collider” in this instance because both *T*\* and *M_d_* cause it. Thus, we see in Figs. 3 and 4 arrows between these random variables, indicative of a possible association, and a square around *I*(*T*\* < *M_d_*), indicative of a conditioning event. Assuming 0 < *P*(*T*\* < *M_d_*) < 1 so the conditioning event is non-trivial, an association between *D* and *T*\* given *G* implies 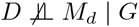, and the converse of this statement is also true. Association between *D* and *T*\* given *G* therefore serves as a powerful and valid proxy for association between *D* and *M_d_* given *G*. These associations result from conditioning on the “collider” *I*(*T*\* < *M_d_*); association paths are opened between *D* and *T*\* given *G*. Were *I*(*T*\* < *M_d_*) not to be conditioned on, *D* and *T*\* would be independent given *G*. The structure of this DAG is identical to that found in classic selection bias (even if the variables are not), where exposure and outcome both cause some indicator that is conditioned on in the analysis, which results in a spurious association between exposure and outcome even under the null.

**Figure 3:**
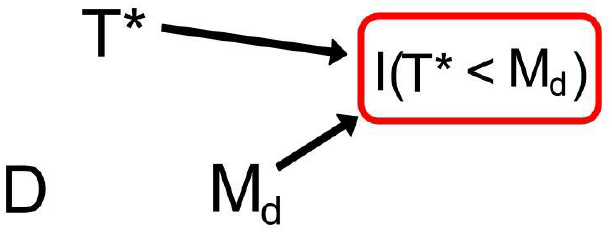
This DAG provides the framework for Test 2. When *D* is not associated with *M_d_*, there is no association between *D* and *T*\*, despite the conditioning event, using rules of DAGs. This figure represents these random variables within each stratum of *G*.

**Figure 4:**
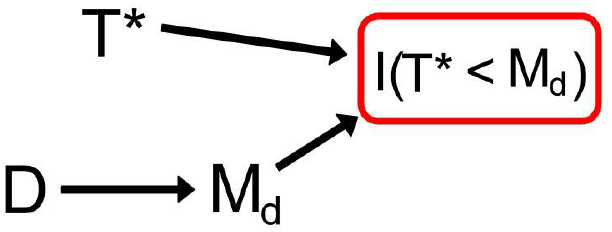
This DAG provides the framework for Test 2. When *D* is associated with *M_d_*, an association between *D* and *T*\* is induced due to the conditioning event using rules of DAGs. This figure represents these random variables within each stratum of *G*.

We could assume *D* and *T*\* are known for all observations in our data set and propose a test of association for these random variables under the framework described above. However, doing so is unrealistic as it assumes follow-up data on age at disease, *D*, for those observed at *T*\* as controls (i.e., those with *D* > *T*\*). Thus, we assume *D* and *T*\* are observed only for cases (i.e., those realizations satisfying *D* < *T*\*) and propose a test of “quasi-independence” between *D* and *T*\* given *G* in the region of *D* < *T*\* given *G*. If we assume that the independence which holds on the region *D* < *T*\* also holds for the entire joint distribution of (*D T*\*)^*T*^, then since there is an association between *D* and *T*\* given *G* if and only if *D* and *M_d_* are associated given *G*, this test is valid. We do not feel this assumption is an overly strong one, but is instead reasonable. In general, a joint distribution that exhibits dependence will not have that structure isolated to a certain region–examination of just a single region will reveal it, even if there are pathological counterexamples where this behavior does not hold.

We describe here this proposed test of quasi-independence. As before, let there be *n* realizations of 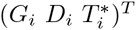, and again define 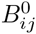 (alternatively, 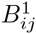) similarly to how we did with 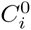 (alternatively, 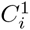), where 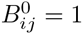 if *G* = 0 and 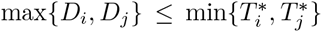 and 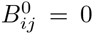 otherwise, and where 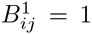 if *G* = 1 and 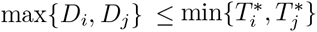 and 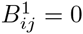 otherwise. Also, define 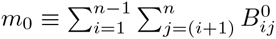 and 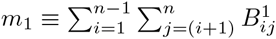.

Then the test statistic for the stratum *G* = *g, W_g_*, with *g* ∈ {0,1}, is

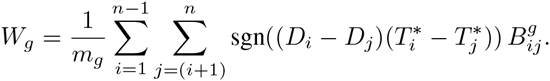

Then *W_g_* ~ *N*(0, *u_g_*), where

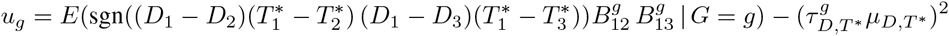

and where 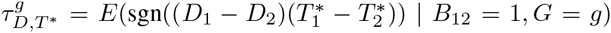 and *μ_D,T*_* = *P*(*B*_12_ = 1). As with Test 1, since we would reject if either *W*_0_ or *W*_1_ falls in some predetermined critical region because dependence between *D* and *M_d_* given either *G* = 0 or *G* = 1 may mean *OR_tr_*(*t*\*) ≠ *OR_ob_*(*t*\*), for a size *α* test, our p-value threshold *α*\* for *W*_0_ and *W*_1_ satisfies *α* = 1 − (1 − α*)^2^. Thus, our test rejects for 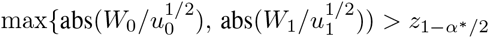.

Power curves for Test 2 as a function of the association between *D* and *M_d_* are shown in Figs. 1 and 2. As with the simulations for Test 1, these curves were generated at 11 different values of Kendall’s *τ*, used as a measure of the association between *D* and *M_d_*. In our case a Kendall’s *τ* value of 0 corresponds to independence between *D* and *M_d_*. The power curve for Test 2 in Fig. 1 was generated using 3000 iterations at each value, while that in Fig. 2 was generated using 1000 iterations at each value, and power was again estimated by averaging over these iterations. In both Figs. 1 and 2, at each iteration the test statistic was calculated using 1000 comparable pairs. *D, T*, and *M_d_* were distributed multivariate normal with means of 5, 9, and 9, respectively, and standard deviations of 0.7. The correlation between *D* and *M_d_* varied as measured by Kendall’s *τ*, while *T* was assumed independent of (*T, M_d_*)^*T*^.

As mentioned at the end of the description of Test 1 and for reasons given there, if this test does not reject *D* ⊥ ⊥ *T*\* | *G*, implying *D* ╨ *M_d_* | *G*, we would again need to stratify the data by *T*\* in order for *OR_ob_*(*t*\*) to be unbiased for *OR_tr_*(*t*\*). In other words, one could split the data in subsets based on age at entry, *T*\*, and calculate stratum-specific odds ratios. It is important to note that the reason for doing so applies no more in the context of testing for Neyman’s bias than it would any standard case-control study, where a mixture of ages at study entry results in an odds ratio that is difficult to interpret and possibly biased as shown in § 4.

### 5.4 Test 3: Estimating population exposure proportion

With knowledge of disease prevalence, we can construct an estimate of the exposure in the general population from case-control study data that is unbiased in the absence of Neyman’s bias, but biased otherwise. Thus, if the exposure proportion in the population is also known, as might be the case in GWAS where minor allele frequencies (MAFs) are oftentimes known for SNPs in different populations, we can test for the presence of Neyman’s bias by examining their discrepancy. We develop one possible hypothesis test below where, again, *H*_0_ is *OR_tr_*(*t*\*) = *OR_ob_*(*t*\*), and *H_a_* is the complement of *H*_0_.

If we make an assumption of *P*(*D* < *M_d_*) = 1, then in comparing *OR_tr_*(*t*\*) and *OR_ob_*(*t*\*), we see that their equivalence depends on

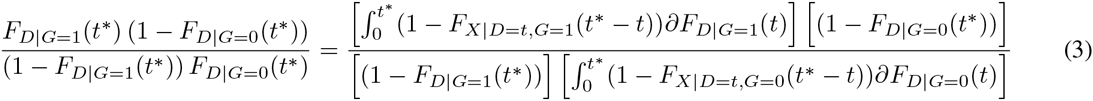

if and only if

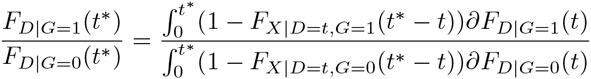

So define *p*_2_(*t*\*) = *P*(*G* = 1|*D* < *t*\*) = *P*(*G* = 1 | Case at *t*\*). Then defining

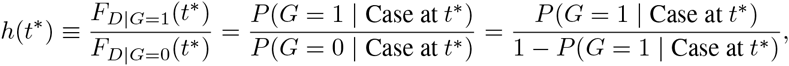

we have *p*_2_(*t*\*) = *h*(*t*\*)/(1 + *h*(*t*\*)), and defining

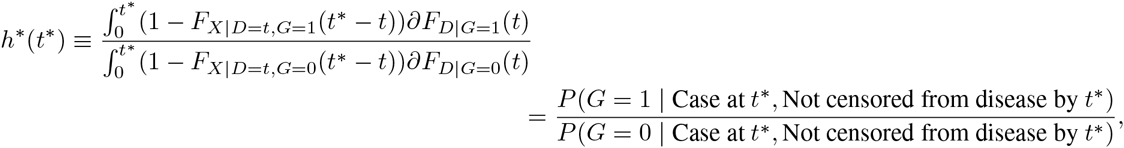

then we have 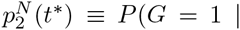 Case at *t*\*, Not censored from disease by *t*\*) = *h*\*(*t*\*)/(1 + *h*\*(*t*\*)). When equation 3 does not hold,

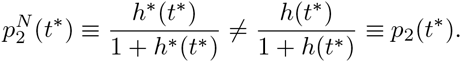

Thus, if bias is present so that *OR_tr_*(*t*\*) ≠ *OR_ob_*(*t*\*), then *h*(*t*\*) ≠ *h*\*(*t*\*), and it will follow that 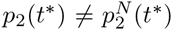. This idea can be leveraged in a hypothesis test if there is external knowledge of the population exposure proportion and population prevalence of disease.

By definition of 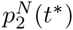, its estimator, 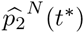, is the observed exposure proportion among cases where 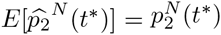. Let *p*_1_(*t*\*) = *P*(*G* =1 | *D* > *t*\*, *M_d_* > *t*\*), and since *P*(*D* < *M_d_*) = 1 by assumption, *p*_1_(*t*\*) = *P*(*G* = 1 | *D* > *t*\*) = *P*(*G* = 1 | Control at *t*\*). Then 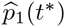 is the observed exposure proportion among controls, and 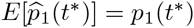.

We will estimate 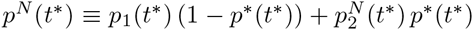 with 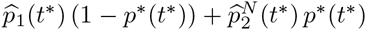. Also, define *p*\*(*t*\*) ≡ *P*(Case at *t*\*) = *P*(*D* < *t*\*), which implies (1 – *p*\*(*t*\*)) = *P*(Control at *t*\*) = *P*(*D* > *t*\*). So *p*\* (*t*\*) is the population prevalence of disease at a common age *t*\* and is considered fixed and known. Since

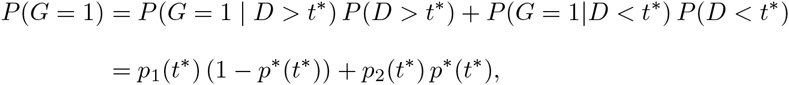

if 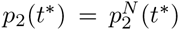, which indicates that *OR_ob_*(*t*\*) = *OR_tr_*(*t*\*), then *p^N^*(*t*\*) = *P*(*G* = 1). Since we consider *P* (*G* = 1) fixed and known, the discrepancy between 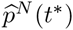 and *P* (*G* = 1) will inform our test.

Define 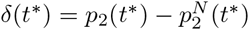. Then

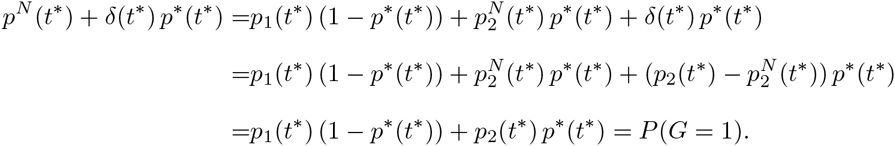

So *P* (*G* = 1) and *p^N^*(*t*\*) differ by *δ*(*t*\*) *p*\*(*t*\*). The variance associated with our estimate of the exposure proportion 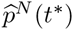 is

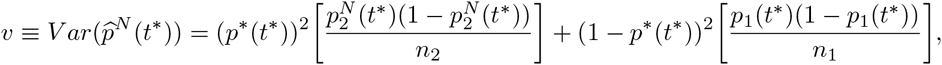

where *n*_2_ is the number of cases and *n*_1_ the number of controls. We can estimate *υ* with 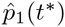 and 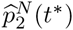 and call the quantity V. So, using a large sample approximation, we can construct an α level hypothesis test for the presence of Neyman’s bias by rejecting for

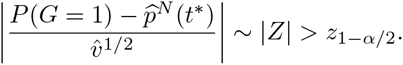

The power becomes

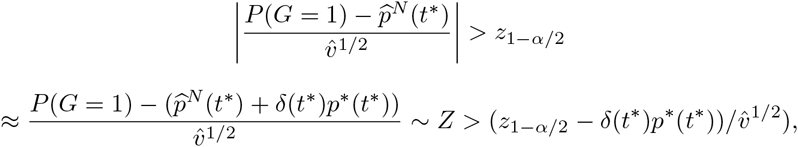

assuming one tail probability negligible. We see that power decreases as *p*\*(*t*\*) decreases and increases with *δ*(*t*\*), interpreted as the “degree of Neyman’s bias.”

Power curves for Test 3 are shown in Fig. 5. Consistent with our understanding of the test, power increases as *p*\*(*t*\*) increases. The type 1 error rate of the test is 0.05. The x-axis of Fig. 5, relative probability of observation, is defined as 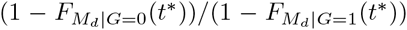; the x-axis begins at one and values greater than one imply that exposed cases are less likely to be sampled than unexposed cases because exposed subjects get disease earlier. Curves were generated using 300 cases and 300 controls in each simulated study, and the population-level exposure proportion was 0.09. Power was calculated based on 4000 iterations at each of the 11 total relative probabilities seen on the x-axis. This entire procedure was done for disease prevalences, *p*\*(*t*\*), of 0.1, 0.2, and 0.3. We assume that there is no variation in *t*\* so that *p*\*(*t*\*) is also fixed.

**Figure 5:**
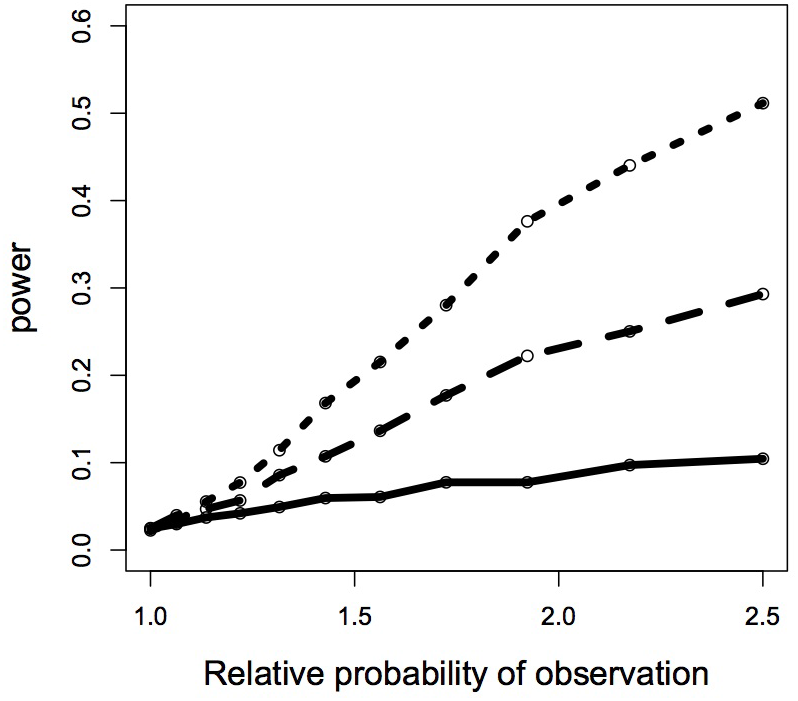
Power for test 3 as a function of *p*\*(*t*\*) and the relative probability of observing the unexposed cases versus exposed cases. As the relative probability increases (i.e., it is more likely to observed unexposed cases than exposed cases) as is the case when there are a greater number of mortality-inducing events among the exposed, there is more bias and power. The solid, dashed, and dotted lines represent population prevalences of disease (*p*\*(*t*\*)) of 0.1, 0.2, and 0.3, respectively.

## Data analysis

### 6.1 Test 1 applied to an atrial fibrillation data set

We applied Test 1 to the Framingham Heart study, a longitudinal study where we would not expect to find evidence of Neyman’s bias; subjects would not be lost to high-mortality diseases by nature of the study design. We would therefore hope that the null hypothesis for Test 1 would not be rejected for each exposure stratum. We consider our exposure to be gender and the disease atrial fibrillation. Among the cohort, there were 82 males and 188 females who had had atrial fibrillation and died of a related cause. Z-statistics calculated using the methodology of Test 1 for the male and female strata are 0.84 and 0.68, respectively, with associated p-values of 0.20 and 0.25. The interpretation of this result is that, were we to calculate an odds ratio for atrial fibrillation at some set age where the exposure is gender, we would not expect to encounter bias. Again, however, the test is used solely for illustration in this case because the study is prospective and not subject to the bias.

### 6.2 Test 2 applied to a brain tumor data set

We apply Test 2 to a brain tumor data set. Seventy-five subjects with oligodendroglioma, a malignant brain tumor, were enrolled in a study at the London Regional Cancer Centre from 1984–1999 (Betensky et al., 2003; Ino et al., 2001). The data set consisted of patient age at diagnosis of oligodendroglioma (i.e., age at disease, D) and age at start of chemotherapy (i.e., entry into the study, *T*\*) in addition to genetic markers and other covariates. We consider the marker at the 1pLOH locus, thought to be potentially associated with tumor sensitivity to chemotherapy. Applying Test 2 to the data set, first within the exposed stratum of the 1pLOH marker, we obtain a Z-statistic of 6.85, significant at the 0.05 level (*p* < 0.001). The sample size was insufficient to apply the test to the unexposed stratum. However, since a significant test statistic within any stratum is sufficient for rejection of the null hypothesis, we reject the null hypothesis of *D* ╨ *M_d_ | G* and conclude that there could be an association between *D* and Md within strata of *G*. The result of the test suggests that if one were to calculate an odds ratio of oligodendroglioma for the 1pLOH marker for subjects at a fixed age, the result might be biased.

Consistent with the comparability criterion of Test 1 being more strict than that for Test 2, the sample size of 75 subjects was insufficient to additionally use Test 1. Had it been used, conclusions drawn from it would not have changed a lack of faith put on the odds ratio in this data set due to results from Test 2.

### 6.3 Test 3 applied to a stroke-mortality data set

We apply Test 3 to a GWAS data set of ischemic stroke coming from a cohort based at Massachusetts General Hospital consisting of 383 cases and 384 controls. We use a wide interval estimate of ischemic stroke prevalence, ranging from 0.5%-5%, based on a search of the stroke literature (Feigin et al., 2009; Johnston et al., 2009; CDC, 2012). With this range of *p*\*(*t*\*), we reconstruct what would be population exposure proportion, which is unbiased for the true population exposure proportion assuming that Neyman’s bias is not present. We calculate a test statistic based on the difference between the true population exposure proportion and our estimate of it, divided by an estimate of the standard error. Using a 0.0005 Bonferroni-adjusted significance level, we find that 42 of the 99 SNPs in the study suggest that Neyman’s bias may be present. The interpretation of this result is that any one of the odds ratios calculated for these 42 SNPs might be biased. We additionally perform a power calculation for this test using realized minor allele frequencies in the data set and generous estimates of both 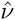, *δ*(*t*\*), and *p*\*(*t*\*). Doing so yields power calculations little above α, at 0.06, which we discuss below.

## Discussion

Test 2 with the brain tumor data suggests that Neyman’s bias may be present because the within exposure stratum association between *D* and *T*\* suggests a within exposure stratum association of *D* and *M_d_*. However, we should restate that an association within strata does not necessary imply that bias is present; it is only when the *D* ╨ *M_d_ | G* holds that we can conclude that Neyman’s bias is not present. Additionally, the study design may contribute to a within stratum association between *D* and *T*\* and so the authors suggest that more work is needed to form stronger conclusions regarding the potential presence of Neyman’s bias in this study.

As with the result from Test 2, the rejection of the null hypothesis of no Neyman’s bias in the stroke-mortality data by Test 3 needs confirmatory analyses. A primary concern is that if the population underlying the measurements in dbSNP, the source of our “true” population MAFs against which we compare the estimate, is significantly different than that composing the study subjects, the type 1 error could be inflated. Since, for many of the SNPs in the data set, the MAF among cases and the MAF among controls did not contain the population MAF, which should be the case as the sample size gets large, there is some evidence of different underlying populations. Another assumption that may not be satisfied is *P*(*D* < *M_d_*) = 1. While *P*(*D* < *M_d_*) = 1 is unlikely to ever be fully satisfied, ischemic stroke is an event with numerous comorbidities and so violations of the assumption may be too large for a valid test (Ostwald et al., 2006; Bots et al., 1997). Lastly, the description of Test 3 showed that the power for detection of bias goes to *α* as the population prevalence of disease gets small. The implication of this result is that any bias detected when the population prevalence of disease ranges over a relatively small 0.5%–5% is more likely due to unsatisfied assumptions than genuine Neyman’s bias. The generous power calculation of 0.06 confirms this belief–it is unlikely that a large proportion of SNPs would have significant p-values, as we have, when there is littler power to detect the bias. It is more likely that the reference population minor allele frequencies are unreflective of the population in the MGH study, which is an assumption that must be satisfied for a valid test.

We did not use Test 1 on the brain tumor and stroke data sets because of an insufficient sample size and insufficient covariates, respectively. The sample size was insufficient in the brain tumor data set because the comparability criterion for Test 1 is more stringent than that for Test 2, so there are only a limited number of pairs of observations that can contribute to estimation of the necessary parameters, especially when overlap between the multivariate random variables (*D T*\* *M_d_*)^*T*^ is minimal. Thus, while Test 2 might be thought of as somewhat removed from testing *D* ╨ *M_d_ | G* because it tests *D* ╨ *T*\* | *G* as a proxy, one advantage of Test 2 over Test 1 is that there are fewer restrictions imposed by the comparability criterion, allowing for more flexible use of the data.

## Acknowledgements and address

The authors wish to thank Dr. Deborah Blacker for many helpful comments used in the preparation of this manuscript as well as Drs. Gregory Cairncross and David Louis for use of the brain tumor data. Dr. Guido Falcone provided invaluable support in the preparation of the ischemic stroke dataset. The MGH ischemic stroke dataset was supported by the American Heart Association/Bugher Foundation Centers for Stroke Prevention Research, the National Institute of Neurological Disorders and Stroke, the Deane Institute for Integrative Study of Atrial Fibrillation and Stroke, and the Keane Stroke Genetics Research Fund. Dr. Anderson is supported by a Clinical Research Training Fellowship from the American Brain Foundation. Prof. Rebecca Betensky is supported by the National Institutes of Health grant CA075971. Dr. Swanson was supported by the National Institutes of Health Training Grant T32 NS048005 while the work was completed at Harvard School of Public Health. His current address is GNS Healthcare, 196 Broadway, Cambridge, MA 02139. His email address is.

## Appendix: direction of bias and examples

We provide a theorem regarding the direction of Neyman’s bias under certain modeling assumptions and examples of when Neyman’s bias does or does not occur.

### Theorem 1

*If G is associated with D such that OR(t*)* ≠ 1, *the distribution of D | (G = 0) and D | (G = 1) belong to the same location family, P (X > 0)* = 1, *P (X < *t*^**^) > 0 (where *t*^**^ is defined as the time between t* and the first possible presence of disease among the exposed or unexposed), and X ╨ (D G)^T^, then OR_ob_(*t*\*) = OR_tr_(t*). Specifically, if D | (G = 0) is stochastically greater than D | (G =1) (alternatively, stochastically less than) so that exposure is a risk factor for disease (alternatively, protective against disease), then OR_ob_(*t*\*) ≠ OR_tr_(t*) (alternatively, OR_ob_(t*) > OR_tr_(t*))*.

### Proof

Define 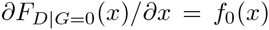 and 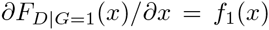, and suppose that *f*_1_(*x*) = *f*_0_(*x − k*) for some *k* positive, without loss of generality. Such a scenario corresponds to exposure being protective against disease, though below we will also consider it a risk factor. *f*_1_(*x*) and *f*_0_(*x*) are in the same location family. Define *F*(*x*) as the cumulative distribution function of *X* evaluated at *x* and remember *F*(0) = 0 and *F*(*t*\*) > 0. Consider the two quantities:

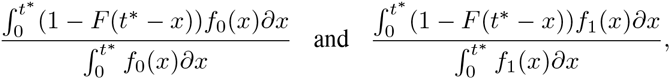

which we call the “percent erosion” of 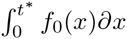 and 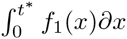, respectively. Then

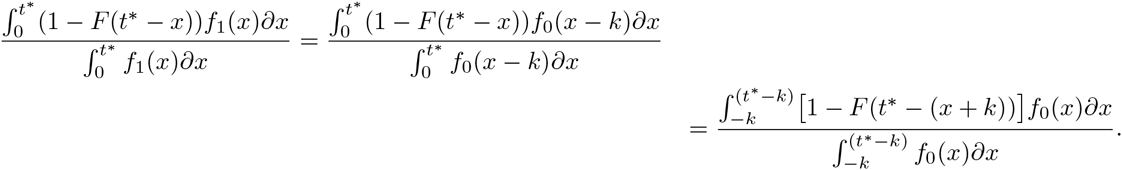

Since *F*(·) a cumulative distribution function and therefore increasing, we have

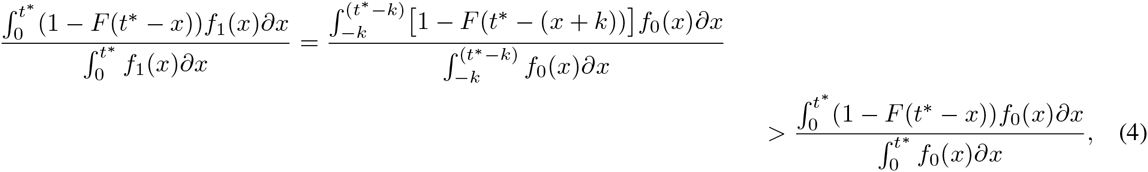

because at every “successive” dx in each integral, 1 − *F*(*t*\* − (*x + k*)) ≥ 1 − *F*(*t*\* − *x*) and there is some 0 < *x* < *t*\* for which 1 − *F*(*t*\* − (*x + k*)) > 1 − *F*(*t*\* − *x*). Thus, the “percent erosion” of *f*_0_(*x*) will always be greater than that of *f*_1_(*x*) = *f*_0_(*x − k*), which is intuitive since *f*_1_(·) is located to the right of *f*_0_(·) and thus subject to the corrosive effects of *F*(·) for less “time.” Then using the inequality in (4),

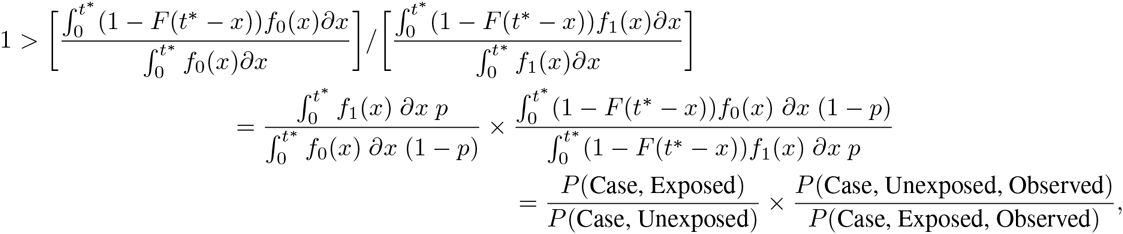

which implies that

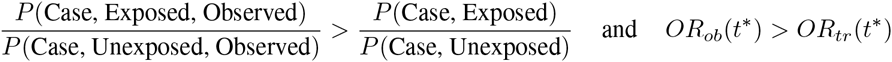

since *P*(*X* > 0) implies *P*(Control, Exposed, Observed) = *P*(Control, Exposed) and *P*(Control, Unexposed, Observed) = *P*(Control, Unexposed). Again, these inequalities only hold when exposure is protective against disease. When exposure is a risk factor for disease and therefore shifts the mean age of disease onset to the left under the above assumptions,

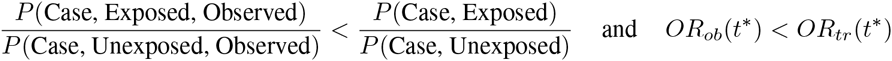

using analogous results. So we see that the bias is not toward the null, but in a definite direction depending on model assumptions.

*Example 1.* Consider *D* | (*G* =1) uniform on (0, 2), *D* | (*G* = 0) uniform on (0,1), and *X* uniform on (0, 3), independent of *G*. Clearly the distributions of disease for exposed and unexposed are not in the same location family in this case, and the model for *X* corresponds to disease-induced mortality necessarily occurring within 3 times units after disease, *D*. We need only consider cases when investigating the odds ratio since we assume *P*(*X* > 0) = 1, implying *P*(*D* < *M_d_*) = 1. Taking *t*\* = 1,

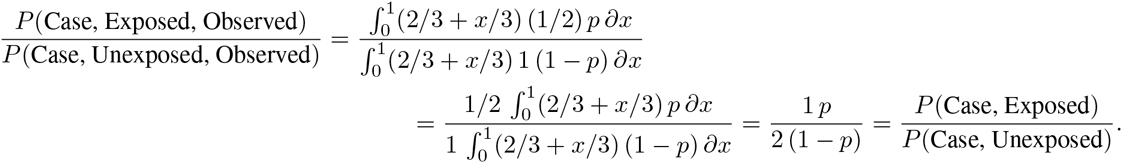

So we have *X* independent of exposure status and time of disease-onset, as was the case above, but here *OR_ob_* = *OR_tr_*.

*Example 2.* Consider again *D* | (*G* = 1) uniform on (0, 2), and *D* | (*G* = 0) uniform on (0,1). However, consider *X* | (*G* = 1) uniform on (0,3) and *X* | (*G* = 0) with density *f_X|G=0_(x)* = 2/3(1 − *x*)^2^ on [0,1 + (9/2)^1/3^]. Again, we need only consider cases when investigating potential bias of the odds ratio since we assume *P*(*D < M_d_*) = 1 so that controls are not subject to the bias-inducing mortality event. Taking *t*\* = 1,

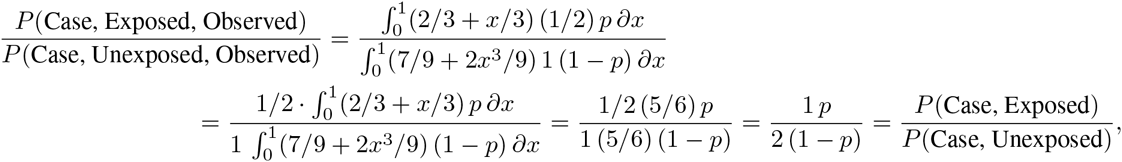

and so here we have no bias again.

*Example 3.* Assume the same models of *D* conditional on *G*, and suppose *X* | (*G* = 1) is uniform on (0,3) and *X* | (*G* = 0) has density *f*_*X|G* = 0_(*x*) = 5/2 (1 − *x*)^4^ on [0,1 + 2^1/5^]. For the reasons given above, we again only consider cases for investigating the bias of the odds ratio. Taking *t*\* = 1,

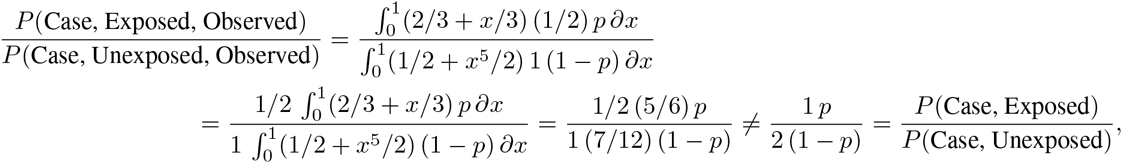

and so here we have bias.

*Example 4.* Take *D* | (*G* =1) with density *f*_*D|G*=1_(*x*) = *x*^2^/4 on [0,12^1/3^], *D* | (*G* = 0) with density *f*_*D*|*G*=0_(*x*) = *x*/3 [0,6^1/2^]. Then let *X* | (*G* = 1) have density *f_X|G=1_*(*x*) = (2−*x*)^2^/4 on [0, 2+4^1/3^] and *X* | (*G* = 0) be uniform on [0,2]. As before, we need only consider cases when investigating the odds ratio since we assume *P*(*D* < *M_d_*) = 1 so that controls are not subject to the bias-inducing mortality event. Taking *t*\* = 2,

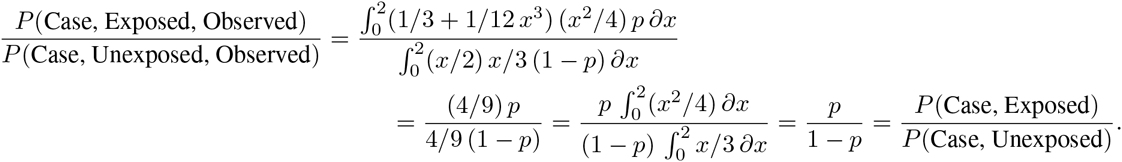

Remember that *P*(Case, Exposed)/*P*(Case, Unexposed) = *p*/(1− *p*) implies *OR_tr_*(*t*\*) = 1 when *P*(*D < M_d_*) = 1, which is assumed from condition 3.

*Example 5.* On the other hand, we can obtain a biased odds ratio using the same conditional disease models as in the previous example and having *X* | (*G* =1) with density *f_X|G=1_*(*x*) = (2 − *x*)^2^/4 on [0,2 + 4^1/3^] and *X* | (*G* = 0) uniform on [0, 2]. We again assume *P*(*D < M_d_*) = 1 from condition 3. Taking *t*\* = 2,

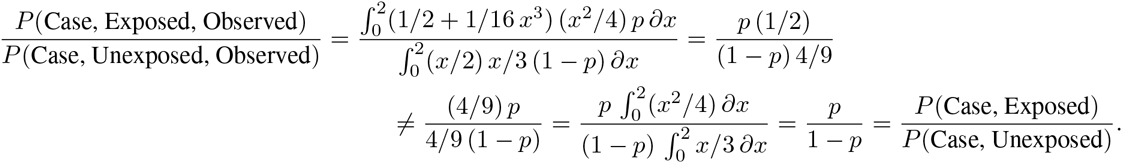

